# Functionalizing lipid sponge droplets with DNA

**DOI:** 10.1101/2021.11.28.470255

**Authors:** Christy Cho, Henrike Niederholtmeyer, Hyeonglim Seo, Ahanjit Bhattacharya, Neal K. Devaraj

**Affiliations:** Department of Chemistry and Biochemistry, University of California, San Diego, USA; Max Planck Institute for Terrestrial Microbiology, Marburg, Germany

**Keywords:** coacervate compartments, DNA, glycolipids, self-assembly, synthetic biology

## Abstract

Nucleic acids are among the most versatile molecules for the construction of biomimetic systems because they can serve as information carriers and programmable construction materials. How nucleic acids interact with membranous coacervate compartments such as lipid sponge droplets is not known. Here we systematically characterize the potential of DNA to functionalize lipid sponge droplets and demonstrate a strong size dependence for sequestration into the sponge phase. Double stranded DNA molecules of more than 300 bp are excluded and form a corona on the surface of droplets they are targeted to. Shorter DNA molecules partition efficiently into the lipid sponge phase and can direct DNA-templated reactions to droplets. We demonstrate repeated capture and release of labeled DNA strands by dynamic hybridization and strand displacement reactions that occur inside droplets. Our system opens new opportunities for DNA-encoded functions in lipid sponge droplets such as cargo control and signaling.

## Introduction

Compartmentalization is an essential property of living cells. Among the many important roles of compartmentalization, it allows cells to regulate reaction rates, maintain reactions in otherwise incompatible conditions, and generate concentration gradients to power important processes like ATP production. Synthetic compartments that self-assemble from simple building blocks in vitro can serve as minimal models for cellular compartments. Coacervates, for example, are molecularly crowded droplets that form by phase separation of macromolecules or amphiphiles from an aqueous solution. Coacervate systems composed of oppositely charged polyions or other interacting macromolecules have been used to emulate the membraneless organelles of eukaryotic cells, such as RNA granules.^[1]^ Being open to the surrounding environment coacervates can spontaneously enrich molecules from the surrounding medium, which has led to enhanced reaction rates for ribozymes,^[2,3]^ transcription^[4]^ and a number of other enzymatic processes.^[5,6]^ As coacervates are held together by non-covalent interactions that can be modulated by environmental conditions, coacervates have been engineered to reversibly assemble and disassemble in response to stimuli such as pH,^[5]^ changes in enzymatic activity,^[1,7–9]^ or temperature.^[10]^ Controlled and reversible sequestration of specific molecules has in turn been used to regulate rates of biochemical reactions that depend on the availability of the sequestered molecules.^[11,12]^ In the process of engineering life-like systems with increasing complexity from the bottom up, synthetic compartments will play an important role in regulating and integrating multiple biochemical functions by locally and dynamically changing the concentrations of the molecules involved in these biochemical reactions. Ideally, compartment-mediated concentration and localization changes should be inducible by signaling molecules, lead to large rapid changes, and have the potential for multiplexing of signals. Additionally, for a rational engineering of novel functions in coacervate compartments, it is essential to perform a systematic characterization of the partitioning of molecular cargo into coacervate droplets. Partitioning depends on properties of the coacervate phase as well as properties of the guest molecules such as charge and molecular weight.^[1,13–15]^

We recently developed lipid sponge droplets, a programmable self-assembled coacervate system, with which we demonstrated acceleration and control of enzymatically catalyzed reactions.^[12]^ Lipid sponge droplets are coacervate compartments that assemble from the glycolipid. *N*-oleoyl β-D-galactopyranosylamine (GOA) and nonionic surfactant octylphenoxypolyethoxyethanol (IGEPAL). Lipid sponge droplets contain dense, nonlamellar lipid bilayer networks and have some similarities to the endoplasmic reticulum. The membranous sponge phase is intersected by nanometric aqueous channels that have characteristic sizes of the order of 4 nm. We demonstrated that droplets doped with small quantities of functionalized lipids could be programmed to efficiently sequester specific affinity-tagged proteins. However, there is a strict threshold due to the size of the water channels, and large protein complexes such as the ClpXP protease were excluded from the lipid sponge phase.

Nucleic acids offer numerous advantages for programming the molecular functions of cell-like structures. Nucleic acids such as DNA can serve as a construction material for structures and machines at the nanometer scale,^[16,17]^ and as an information carrier for the synthesis of proteins,^[18]^ or DNA-based computing.^[19]^ Dynamic DNA nanotechnology leverages the high degree of programmability and predictability of DNA base pairing rules to construct dynamic circuits. DNA-based computations often take advantage of toehold mediated strand displacement reactions, where short single-stranded extensions of DNA duplexes help to initiate DNA-branch migration reactions.^[20]^ Toehold mediated strand displacement reactions have been used for information processing and signaling^[21,22]^ as well as the control of material properties.^[23,24]^ It is still unknown how nucleic acids interact with lipid sponge droplets, to what extent they can be sequestered, and if they can be used to program droplet functions. Here, we develop an efficient strategy to target specific nucleic acids into lipid sponge droplets and use double-stranded DNA as a molecular ruler to determine the size cut-off for sequestration into the droplet phase. These findings provide the basis for programming lipid sponge droplets with DNA-encoded functions. As an example, we show repeated sequestration and release of molecular cargo driven by a toehold mediated strand displacement reaction. Our results demonstrate how control of dynamic localization of molecular cargo can be achieved in DNA functionalized droplets, thus offering a new approach to regulate timing and sequence of biochemical reactions.

## Results and discussion

To systematically characterize the sequestration of macromolecules as a function of size, we prepared a series of double-stranded DNA constructs between 28 and 959 base pairs in length each carrying a biotin and a TYE665 fluorophore modification on the 5’ ends (**Supplementary Fig. 1**). To target these DNA constructs into lipid sponge droplets we preincubated the DNA with streptavidin and subsequently mixed the DNA-streptavidin complexes with lipid sponge droplets that were doped with small amounts of biotinyl PE as previously described for protein sequestration^[12]^ (**Fig. 1**). Without the full targeting system, DNA did not partition into the droplet phase (**Supplementary Fig. 2**).

**Figure 1.**
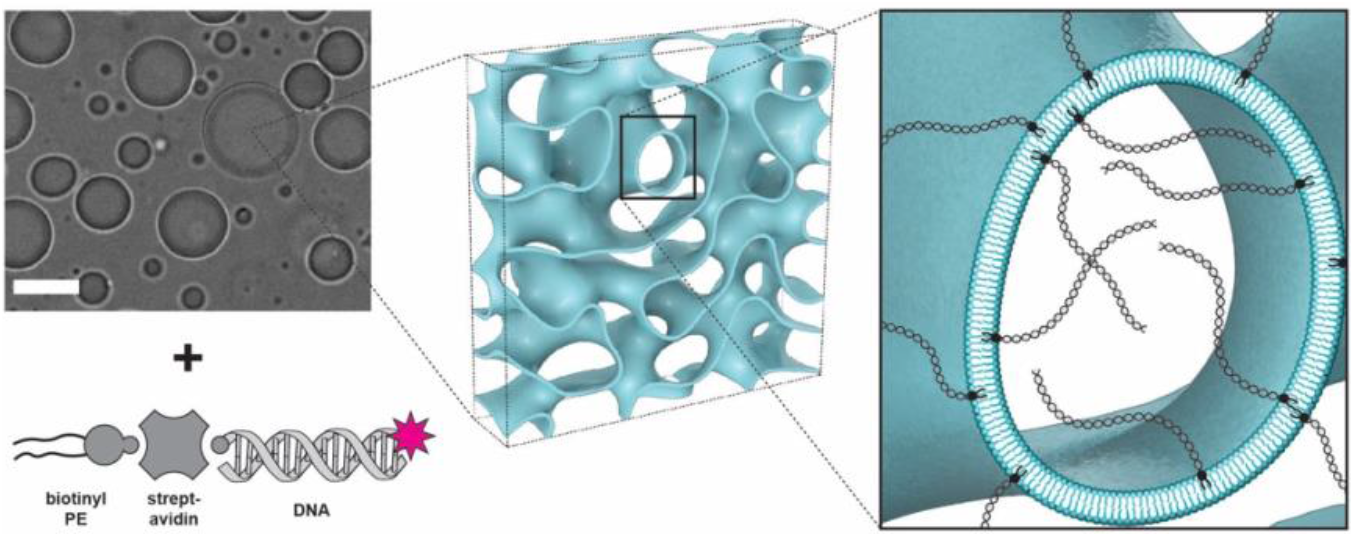
Programming the sequestration of DNA by lipid sponge droplets. Double-stranded DNA constructs of different lengths containing a biotin and a fluorophore modification on the 5’ ends are produced by PCR. DNA is targeted into lipid sponge droplets doped with biotinyl PE through interaction with streptavidin. Scale bar: 25 μm.

In the presence of all targeting components, efficient partitioning of DNA constructs up to 93 bp in length into the droplet phase could be observed by confocal microscopy (**Fig. 2A**). DNA constructs between 108 bp and 208 bp partitioned into the droplet phase with decreasing efficiency, and DNA constructs of 308 bp and larger were completely excluded from the droplet phase. Excluded DNA constructs formed a corona with increased brightness observed around the edges of the droplets (**Fig. 2A, B**). For DNA constructs of intermediate lengths between 108 bp and 158 bp, droplet fluorescence was homogeneous immediately after mixing but, following an incubation for 1-2 h, fluorescence increased around the edges. The phenomenon was particularly prominent in smaller droplets (**Supplementary Fig. 3**). This observation indicates that for DNA constructs of 108 bp and larger, the diameter of the aqueous channels in the droplet phase is a constraint, which leads to the gradual accumulation of DNA constructs at the solution-droplet interface. The partition coefficient is a measure for the enrichment in concentration of a molecule in the droplet phase relative to the solution phase. Here, after correcting for background fluorescence, the partition coefficient was calculated as the ratio between fluorescence in the center of a droplet to fluorescence in solution immediately after mixing DNA with droplets (**Fig. 2C**). Partition coefficients of DNA constructs up to 93 bp in length were around 20, while for DNA constructs with 308 bp and longer slightly negative partition coefficients were calculated, indicating exclusion of DNA constructs from the droplet interior. DNA molecules between 108 bp and 208 bp in length were classified as partially excluded based on their gradually decreasing partition coefficients and the appearance of a corona around the droplets, which appeared faster and became more pronounced with increasing DNA length (**Fig. 2A, Supplementary Fig. 3**).

**Figure 2.**
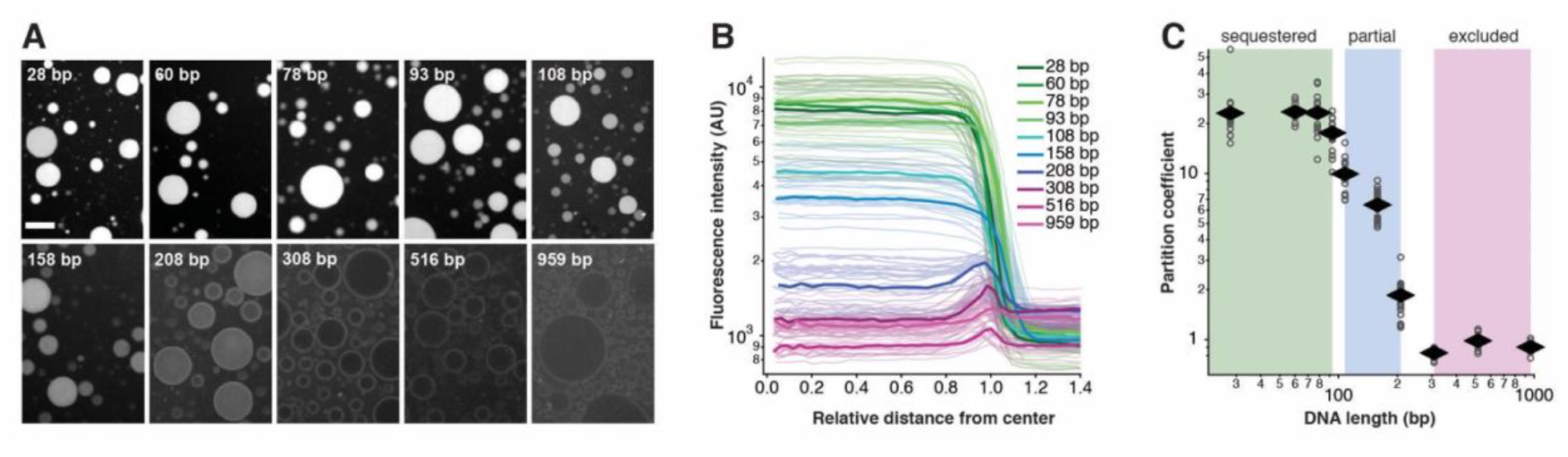
Transition from sequestration to exclusion of DNA targeted into lipid sponge droplets. (**A**) Representative fluorescence images of droplets after mixing with DNA. Images are shown at the same scale, with the same brightness settings in a logarithmic scale for better visualization. Scale bar: 25 μm. (**B**) Fluorescence intensity profiles of droplets. A relative distance of 1 corresponds to the droplet edge and 0 is the center. Thin lines show individual measurements of fluorescence profiles from multiple images and at least 9 droplets per DNA length. Thick lines show a representative profile for each DNA length for better visualization. (**C**) Partition coefficients calculated from background subtracted intensity profiles in (B) (see Methods). Diamond symbols represent the average and open circles the individual values.

The average channel diameter of lipid sponge droplets was previously determined by small-angle X-ray scattering to be 4.1 nm, and we have observed the exclusion of ClpXP protease, a 14.5 nm by 11 nm protein complex.^[12]^ Based on the analysis of Son et al.^[25]^ we estimate the average length of a 93 bp long dsDNA molecule that is anchored to a lipid bilayer at one end to be about 15 nm. We therefore conclude that DNA molecules with a length of almost four times the average channel diameter can be sequestered into the sponge phase. In contrast to the ClpXP protease, linear dsDNA extends primarily in a single dimension. With their small width, DNA molecules can probably be accommodated in the tortuous channel network more easily than most protein complexes that extend in three dimensions. The dynamic and fluid nature of the sponge phase explains the slow expulsion of DNA with lengths of 108 to 208 bp, compared to longer DNA molecules that were completely and immediately excluded from the sponge phase.

Efficient and fast sequestration of short dsDNA molecules into lipid sponge droplets suggested that cycles of uptake and release of molecular cargo could potentially be driven by dynamic DNA networks. We adapted previous designs of toehold mediated strand displacement reactions for controlled capture and release of DNA strands by lipid sponge droplets.^[20,23,26]^ Our strategy (**Fig. 3A**) uses a biotinylated single-stranded DNA capture strand (C_s_) that is anchored in lipid sponge droplets via streptavidin and biotinyl PE. A partially complementary and fluorescently labeled DNA strand F gets sequestered into droplets by base paring with C_s_ to form a partially double-stranded complex C_d_. In addition to the complementary region, strand F has a toehold domain that remains single stranded. When F*, the complementary strand to F, is added, it displaces F from the capture strand. F and F* then leave the droplet as the double-stranded waste product W. This leaves the capture strand in its initial single stranded state. By sequentially adding equimolar amounts of F and F* we should be able to cycle droplets through multiple rounds of binding and unbinding events on demand (**Fig. 3B**). To distinguish each hybridization step during the exchange of cargo we used two F strands that were labelled with different fluorophores. F_red_ carried the TYE665 fluorophore and F_green_ was modified with ATTO488.

**Figure 3.**
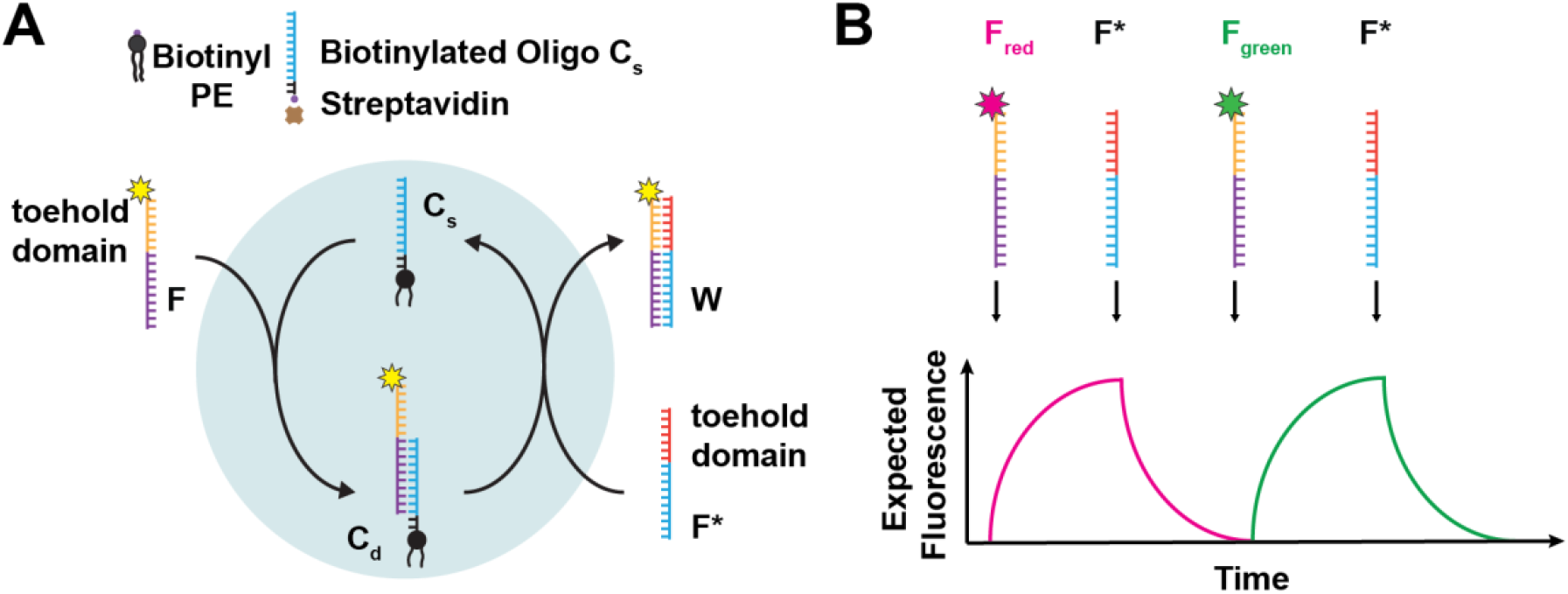
Design of a toehold-mediated DNA strand displacement reaction driving cargo exchange in droplets. (**A**) Schematic of hybridization and toehold mediated strand displacement reactions driving sequestration and release of DNA strands by lipid sponge droplets. Single-stranded capture strand C_s_ is anchored in droplets via a biotin-streptavidin interaction and helps sequester a fluorescently labelled cargo strand F by hybridization forming partially double stranded complex C_d_. The cargo can be removed from droplets by the addition of F* that displaces F from the capture strand by toehold mediated strand displacement. Subsequently, F and F* leave the droplet as double stranded waste product W, leaving the capture strand C_s_ in its initial single-stranded form, ready to capture another cargo. (**B**) Expected changes in fluorescence intensities of sponge droplets in response to the addition of fluorescently labeled oligos F_red_ or F_green_, which can be displaced from droplets by their complementary sequence F*.

To assess the functionality of our strategy, we followed the sequential hybridization and subsequent displacement steps via confocal microscopy (**Fig. 4A**). C_s_ was anchored in droplets via biotin-streptavidin interactions. The sample of droplets was then placed into a well of a chamber slide for timelapse imaging at 37 °C. Initially, at time 0 h, the droplet sample only contained single stranded capture strand C_s_ at a concentration 40 nM. Consequently, droplets showed no fluorescence. When at 0.1 h, complementary strand F_red_ was added in an equimolar amount to C_s_, we observed an immediate increase of fluorescence in droplets (**Fig. 4A**). Fast hybridization of F_red_ forming duplex C_d_ in droplets was also apparent in the fluorescence traces that were extracted from selected regions of the imaged sample (**Fig. 4B**). In a control experiment we verified that the rapidly increasing TYE665 fluorescence was indeed due to specific interactions between the immobilized capture strand C_s_ and F_red_. Droplets without the bound capture strand did not increase in fluorescence, indicating that the sequestration of F_red_ was due to specific base pairing between the complementary strands in the droplets (**Supplementary Fig. 4**).

**Figure 4.**
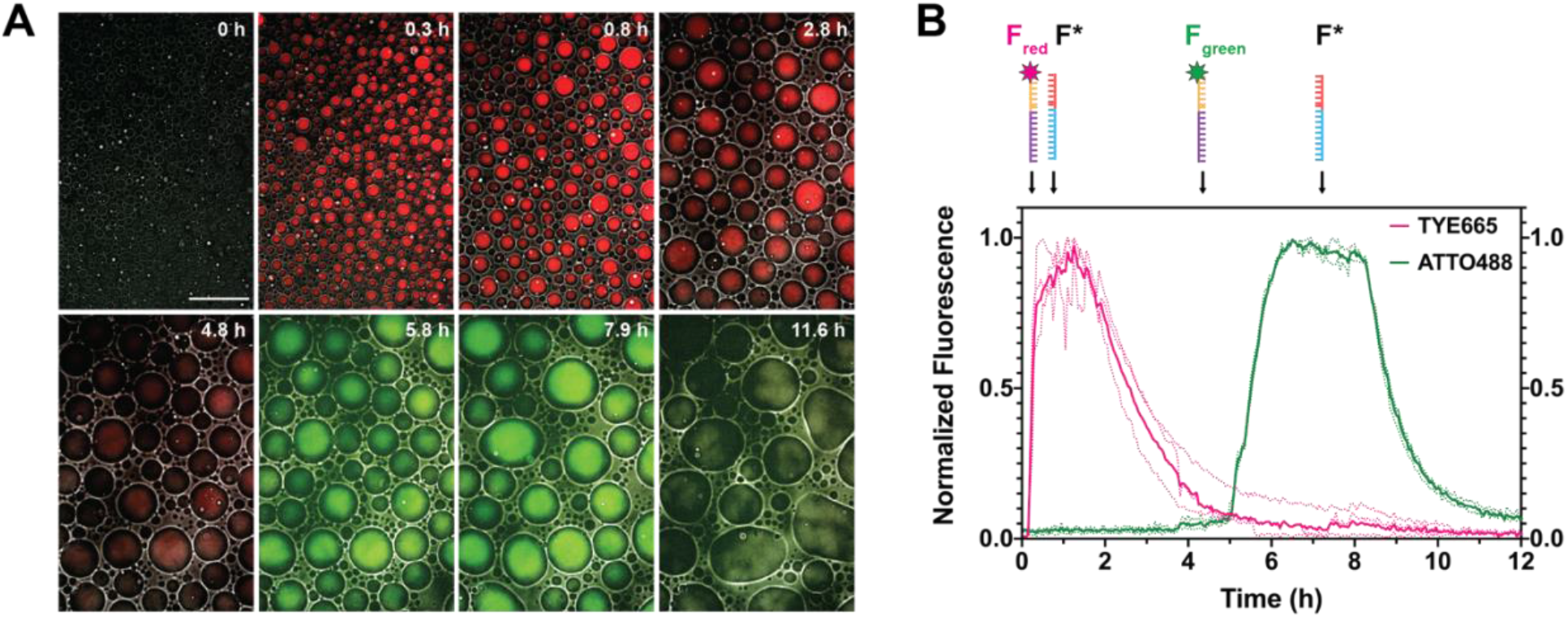
Completion of two cycles of DNA-driven cargo exchange in lipid sponge droplets. (**A**) Confocal microscopy images of sequential hybridization and displacement reactions in lipid sponge droplets. Selected images from time lapse experiment with merged brightfield, TYE665 (red) and ATTO488 (green) channels. Image intensities are window leveled to the same values. Scale bar: 100 μm. (**B**) Kinetics of the change in fluorescence intensities of sponge droplets during the hybridization and DNA displacement reactions. Normalized fluorescence intensities of three selected individual droplets located in different positions in the reaction chamber (dotted lines), and the average (bold) in corresponding channels. F_red_, F*, F_green_ and F* were added at 0.1, 0.8, 4.8, and 7.9 h respectively.

A decreasing rate of fluorescence increase indicated that hybridization of F_red_ in droplets was approaching saturation. F_red_ was then released from droplets by adding an equimolar amount of the invading strand F*, which was designed to form the double-stranded waste product W after hybridizing to F_red_ in a toehold mediated strand displacement reaction. Strand displacement and release of F_red_ could be observed after a short delay by decreasing droplet fluorescence. Even though fluorescence of F_red_ was not quenched, dilution of the released waste duplex W in the solution leads to decreasing fluorescence in the droplets.

When droplet fluorescence approached background levels, we initiated a second cycle of cargo uptake, this time adding a different cargo strand, F_green_, which carried the ATTO488 fluorescent label. Again, after a short delay, hybridization and sequestration into droplets was rapid. When fluorescence plateaued, F* was added. Again, following a delay, droplet fluorescence started decaying, demonstrating the completion of two successful cycles of on-demand cargo uptake and release (**Supplementary Movie 1**).

We observed a difference in the kinetics for the hybridization reaction (sequestration of F_red_ and F_green_ into droplets) compared to the toehold mediated strand displacement reaction (release of F_red_ and F_green_ from droplets). Sequestration was much more rapid than the release, which can be explained by the fact that hybridization between two single-stranded oligonucleotides is very fast. In contrast, release of the fluorescent cargo F required hybridization of F* to the toehold on F and a subsequent three-way branch migration reaction, which is a slower process.^[20]^ The fluorescence release from droplets could be sped up by doubling the molar concentrations of the invading strand F* to two equivalents of the F_red_ concentration (**Supplementary Fig. 5**). The delays in the reaction, between adding a DNA strand and the beginning of a change in droplet fluorescence, were likely due to diffusion of the added strands through the chamber and the molecularly crowded sponge phase of droplets, where diffusion is slowed.^[12]^ Despite the observed delays, reactions completed at comparable time scales to toehold mediated strand displacement reactions in well mixed solutions.^[26]^ Our results demonstrate that toehold mediated strand displacement reactions are a viable approach to control capture and release of molecular cargo by lipid sponge droplets on demand. DNA-based localization control led to higher fold-changes in cargo loading and release than a light-controlled protein-protein interaction^[12]^ and DNA base pairing interactions have the advantage that they can be programmed and multiplexed easily.

In conclusion, we have systematically characterized the potential of DNA functionalization in lipid sponge droplets. Our results lay the foundation for DNA-templated chemistry in lipid sponge droplets and at their solution interface. Double stranded DNA constructs that are too long to be accommodated in the nanometric channels of the lipid sponge phase accumulate on droplet surfaces. Length-dependent surface decoration of droplets with DNA could be used to program physical interactions between individual droplets or droplets and substrates in the future. In contrast, double stranded DNA up to approximately 100 bp in length is sequestered into the interior of lipid sponge droplets with high efficiency. While this length is too short to encode proteins, RNA signals^[27]^ could potentially be produced by transcription in droplets. Here we demonstrate that we can take advantage of the open nature of the lipid sponge phase and use DNA strands as signals to initiate hybridization and strand displacement reactions in droplets. These reactions control uptake and release of molecular cargo that is coupled to DNA. In the future, the system could be extended to include communication between individual neighboring droplets through nucleic acid signals that could be released locally upon activation.^[28–30]^ Communication and computations based on nucleic acids as signals can be easily programmed and multiplexed opening up new opportunities for molecular localization control and signal processing in lipid sponge droplets.

## Acknowledgements

This work was supported by the NSF (CHE-1844346) and the Department of Defense (Army Research Office) through the Multidisciplinary University Research Initiative (Award W911-NF-13-1-0383). HN acknowledges funding by the Deutsche Forschungsgemeinschaft (NI 2040/1-1). We thank Veronica Falconieri Hays for illustration of the lipid sponge phase.

## References

[1] W. M. Aumiller, C. D. Keating, Nat. Chem. 2016, 8, 129–137.

[2] R. Poudyal, R. M. Guth-Metzler, A. J. Veenis, E. A. Frankel, C. D. Keating, P. C. Bevilacqua, Nat. Commun. 2019, 10, 490–13.

[3] K. Le Vay, E. Y. Song, B. Ghosh, T. Y. D. Tang, H. Mutschler, Angew Chem Int Ed Engl 2021, DOI 10.1002/anie.202109267.

[4] E. Sokolova, E. Spruijt, M. M. K. Hansen, E. Dubuc, J. Groen, V. Chokkalingam, A. Piruska, H. A. Heus, W. T. S. Huck, Proc. Natl Acad. Sci. USA 2013, 110, 11692–11697.

[5] S. Koga, D. S. Williams, A. W. Perriman, S. Mann, Nat. Chem. 2011, 3, 720–724.

[6] T. Beneyton, C. Love, M. Girault, T. Y. D. Tang, J.-C. Baret, ChemSystemsChem 2020, 148, 3–9.

[7] S. N. Semenov, A. S. Y. Wong, R. M. van der Made, S. G. J. Postma, J. Groen, H. W. H. van Roekel, T. F. A. de Greef, W. T. S. Huck, Nat. Chem. 2015, 7, 160–165.

[8] K. K. Nakashima, J. F. Baaij, E. Spruijt, Soft Matter 2018, 14, 361–367.

[9] B. S. Schuster, E. H. Reed, R. Parthasarathy, C. N. Jahnke, R. M. Caldwell, J. G. Bermudez, H. Ramage, M. C. Good, D. A. Hammer, Nat. Commun. 2018, 9, eaaf4382–12.

[10] N.-N. Deng, W. T. S. Huck, Angew. Chem. Int. Ed. 2017, 56, 9736–9740.

[11] J. R. Simon, S. A. Eghtesadi, M. Dzuricky, L. You, A. Chilkoti, Mol. Cell 2019, 75, 66–75.e5.

[12] A. Bhattacharya, H. Niederholtmeyer, K. A. Podolsky, R. Bhattacharya, J.-J. Song, R. J. Brea, C.-H. Tsai, S. K. Sinha, N. K. Devaraj, Proc. Natl Acad. Sci. USA 2020, 3, 202004408.

[13] T. J. Nott, T. D. Craggs, A. J. Baldwin, Nat. Chem. 2016, 8, 569–575.

[14] K. A. Black, D. Priftis, S. L. Perry, J. Yip, W. Y. Byun, M. Tirrell, ACS Macro Lett. 2014, 3, 1088–1091.

[15] E. A. Frankel, P. C. Bevilacqua, C. D. Keating, Langmuir 2016, 32, 2041–2049.

[16] K. Jahnke, M. Weiss, C. Frey, S. Antona, J.-W. Janiesch, I. Platzman, K. Göpfrich, J. P. Spatz, Adv. Funct. Mater. 2019, 29, 1808647–8.

[17] B. Yurke, A. J. Turberfield, A. P. Mills, F. C. Simmel, J. L. Neumann, Nature 2000, 406, 605–608.

[18] H. Niederholtmeyer, C. Chaggan, N. K. Devaraj, Nat. Commun. 2018, 9, 5027.

[19] A. Joesaar, S. Yang, B. Bögels, A. van der Linden, P. Pieters, B. V. V. S. P. Kumar, N. Dalchau, A. Phillips, S. Mann, T. F. A. de Greef, Nat. Nanotechnol. 2019, 14, 369–378.

[20] F. C. Simmel, B. Yurke, H. R. Singh, Chem. Rev. 2019, 119, 6326–6369.

[21] H. Liu, Q. Yang, R. Peng, H. Kuai, Y. Lyu, X. Pan, Q. Liu, W. Tan, J Am Chem Soc 2019, 141, 6458–6461.

[22] E. Magdalena Estirado, A. F. Mason, M. Á. Alemán García, J. C. M. van Hest, L. Brunsveld, J Am Chem Soc 2020, 142, 9106–9111.

[23] R. Hager, A. Arnold, E. Sevcsik, G. J. Schütz, S. Howorka, Langmuir 2018, 34, 15021–15027.

[24] A. Buchberger, H. Saini, K. R. Eliato, A. Zare, R. Merkley, Y. Xu, J. Bernal, R. Ros, M. Nikkhah, N. Stephanopoulos, ChemBioChem 2021, 22, 1755–1760.

[25] S. Son, S. C. Takatori, B. Belardi, M. Podolski, M. H. Bakalar, D. A. Fletcher, Proc. Natl Acad. Sci. USA 2020, 117, 14209–14219.

[26] D. Y. Zhang, E. Winfree, J Am Chem Soc 2009, 131, 17303–17314.

[27] M. Weitz, J. Kim, K. Kapsner, E. Winfree, E. Franco, F. C. Simmel, Nat. Chem. 2014, 6, 295–302.

[28] G. Gines, A. S. Zadorin, J. C. Galas, T. Fujii, A. Estévez-Torres, Y. Rondelez, Nat. Nanotechnol. 2017, 12, 351–359.

[29] N. Martin, L. Tian, D. Spencer, A. Coutable-Pennarun, J. L. R. Anderson, S. Mann, Angew. Chem. 2019, 131, 14736–14740.

[30] S. Yang, P. A. Pieters, A. Joesaar, B. W. A. Bögels, R. Brouwers, I. Myrgorodska, S. Mann, T. F. A. de Greef, ACS Nano 2020, 14, 15992–16002.

