## Supporting Information for "Functionalizing lipid sponge droplets with DNA"

#### Methods

##### DNA constructs

Double-stranded DNA constructs were assembled using primers listed in Supplementary Table 1. The 28 bp DNA was assembled by annealing two complementary DNA strands. PCRs were carried out using the NEB Q5 HotStart Master mix in reactions containing 6% (v/v) DMSO using pET-Trx-mSa2 as a template. pET-Trx-mSA2 was a gift from Sheldon Park (Addgene plasmid # 52320; http://n2t.net/addgene:52320; RRID:Addgene_52320). Constructs of 108 bp and larger were assembled in a two-step PCR protocol, where the product of the first PCR served as the template for amplification with the final primers containing the biotin and TYE665 modifications. As necessary to obtain a pure final product, the products of the first PCR reaction were purified using the PureLink™ PCR Purification Kit (Thermo Fisher Scientific) using the supplied high-cutoff buffer or by performing a stab of the correct band on an agarose gel. Final PCR products were checked for purity by agarose gel electrophoresis imaging for GelGreen stain and the TYE665 fluorophore using a Typhoon FLA 9500 laser scanner. Note that the TYE665 fluorophore was better suited for assessing purity because it is more sensitive for small DNA products that could cause false positive sequestration results. Final DNA constructs were purified using the PureLink™ PCR Purification Kit (Thermo Fisher Scientific) using the supplied standard buffer. Concentrations were determined photometrically using a NanoDrop.
DNA oligonucleotides for hybridization and strand displacement reactions in droplets were ordered HPLC-purified and with the modifications listed in Supplementary Table 2. All oligonucleotides used in this study were ordered from IDT.

##### Droplet formation

GOA was synthesized as described previously.[1] IGEPAL was purchased from Sigma (product number I8896, Lot MKCC9036). Biotinyl PE doped droplets were prepared by hydrating GOA and biotinyl PE (Avanti) with IGEPAL at a molar ratio of 1:0.025:0.75 (GOA:biotinyl PE:IGEPAL) in 0.1 M HEPES ((4-(2-hydroxyethyl)-1-piperazineethanesulfonic acid) buffer solution, pH 8.0 at room temperature with the following protocol:

In a glass vial a lipid film was formed from:
50 µL chloroform
10 µL 10 mM GOA
1 µL 2.5 mM biotinyl PE

The lipid film was rehydrated by vortex mixing with 50 µL of rehydration solution:
42.5 µL 100 mM HEPES pH 8.0
7.5 µL 10 mM IGEPAL

**DNA sequestration, hybridization and toehold mediated strand displacement experiments**
For DNA sequestration into droplets, biotinylated DNA (double-stranded constructs or oligonucleotide $C_s$) was mixed with streptavidin at 80 nM DNA and 2 µM streptavidin and incubated for 5 min at room temperature for binding. The DNA-streptavidin solution was then mixed with an equal volume of biotinyl PE droplets by pipetting up and down carefully to allow for binding of the streptavidin-DNA complexes to droplets. Final concentrations in the samples were 40 nM DNA, 1 µM streptavidin, 25 µM biotinyl PE, 1 mM GOA and 0.75 mM IGEPAL. Samples with double stranded DNA constructs were prepared in 5 µL volumes and imaged sandwiched between cover glass.
For hybridization and toehold mediated strand displacement reactions, we prepared 100 µL droplet samples, which were transferred to an 8-well Lab-Tek chamber slide (Sigma Aldrich) and maintained at 37 ºC, 5% $CO_2$ using a stagetop incubator (Okolab, Italy). To the droplet sample solution, equimolar concentrations in respect to $C_s$ (40 nM) of strands F then F* to $C_s$ were added along the wall to avoid disturbing the droplets. To achieve hybridization, 40 nM of fluorescently labeled strand F ($F_{red}$ or $F_{green}$) was incubated with the droplet samples containing $C_s$. After complete saturation of strand F, the solution was incubated with complementary strand F* (40 nM) for toehold mediated displacement of F. Strand $F_{red}$ was added at 0.1 h, F* at 0.8 h, $F_{green}$ at 4.8 h, and the final F* at 7.9 h. Hybridization and displacement reactions were followed by confocal microscopy time lapse imaging acquired every 3 minutes.

**Imaging and image analysis**
Microscopy images were acquired on a Yokagawa spinning-disk system (Yokagawa, Japan) built around an Axio Observer Z1 motorized inverted microscope (Carl Zeiss Microscopy GmbH, Germany) with a 63x, 1.40 NA oil immersion objective or 20x 0.8 NA objective with an ORCA-Flash4.0 V2 Digital CMOS camera (Hamamatsu, Japan) using ZEN Blue imaging software (Carl Zeiss Microscopy GmbH, Germany). The fluorophores were excited with diode lasers (488 nm-30 mW, and 638 nm-75 mW).
For the quantitative analysis of DNA sequestration images were acquired with the 20x objective. Images were analyzed in Fiji/ImageJ[2] using the Radial Profile Extended plugin to obtain smoothed average intensity profiles of droplets. We analyzed circular droplets that had a radius between 9 µm and 24 µm. The integration radius was increased by a factor of 1.5x to include the surrounding solution. The starting and integration angels were chosen to exclude other droplets nearby to get an accurate value of the fluorescence of the solution (**Supplementary Fig. 6**). The distance to center values were then normalized by the droplet radius. For each analyzed droplet we calculated the partition coefficient. For this we used the average of the first 3 values in the

center and the average of the final 3 values in solution from radial profiles, subtracted the background Cy5 fluorescence of a droplet-water sample and calculated the ratio of internal to external concentration.

In cargo-exchange experiments, time lapse images of DNA hybridization and displacement in sponge droplets were analyzed using Fiji/Image J to quantify the average fluorescence intensities of droplets in the TYE665 and ATTO488 fluorescence channels. Droplet fluorescence intensities over time, $F_{droplet}$, were extracted from regions of interest (ROIs) inside individual droplets. Due to droplet coalescence, ROIs were carefully selected for locations that were occupied by a single droplet over the course of the entire experiment. For both fluorescence channels, droplet fluorescence intensities ($F_{droplet}$) over time (t) were normalized:

$$\frac{F_{droplet}(t) - F_{min}}{F_{max} - F_{solution}(t)}$$

$F_{max}$ and $F_{min}$ are the maximum and minimum intensity value of the $F_{droplet}$ values over time. $F_{solution}$ is the fluorescence intensity of a region in the movie that was never occupied by a droplet. The normalized fluorescence intensities of ROIs in three individual droplets were then averaged (Figure 4B).

**Supplementary Movie 1**

**Hybridization and toehold mediated strand displacement reactions in lipid sponge droplets.** Confocal microscopy images of sequential hybridization and displacement reactions in lipid sponge droplets. Selected images from time lapse experiment with merged brightfield, TYE665 (red) and ATTO488 (green) channels. This video corresponds to the images shown in Figure 4.

### References


[1] A. Bhattacharya, H. Niederholtmeyer, K. A. Podolsky, R. Bhattacharya, J.-J. Song, R. J. Brea, C.-H. Tsai, S. K. Sinha, N. K. Devaraj, *Proc. Natl Acad. Sci. USA* **2020**, *3,* 202004408.

[2] J. Schindelin, I. Arganda-Carreras, E. Frise, V. Kaynig, M. Longair, T. Pietzsch, S. Preibisch, C. Rueden, S. Saalfeld, B. Schmid, et al., *Nat. Meth.* **2012**, *9*, 676–682.

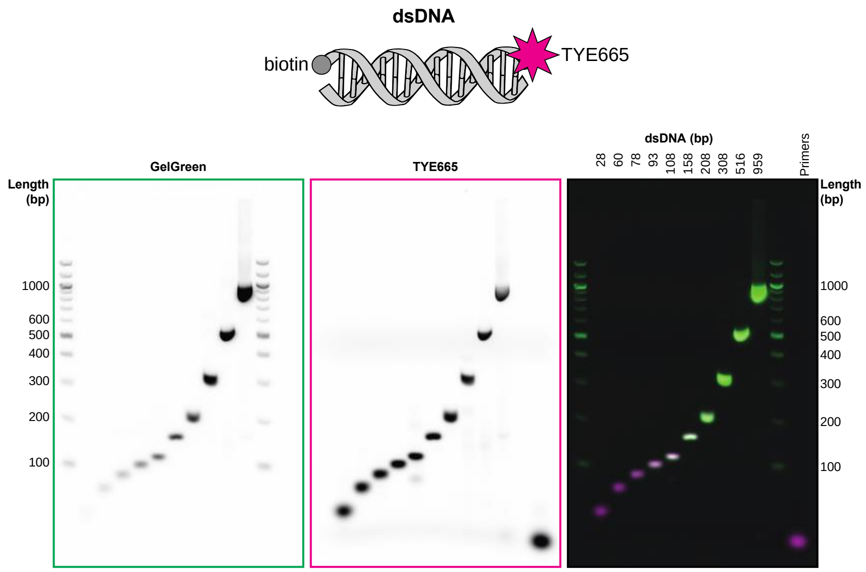


**Supplementary Fig. 1. Agarose gel analysis of DNA constructs.** Gel was 3% agarose in TAE buffer. Per lane 0.5 pmol DNA were loaded (1µl of 500 nM). Lanes contain DNA constructs of the indicated sizes. Control lane labeled as “Primers” contains of 0.5 pmol of each final primer. The gel was stained with GelGreen nucleic acid dye and imaged in the green and Cy5 fluorescence channel on a Typhoon scanner. Channels are shown individually as indicated and as a merged fluorescence image (right).

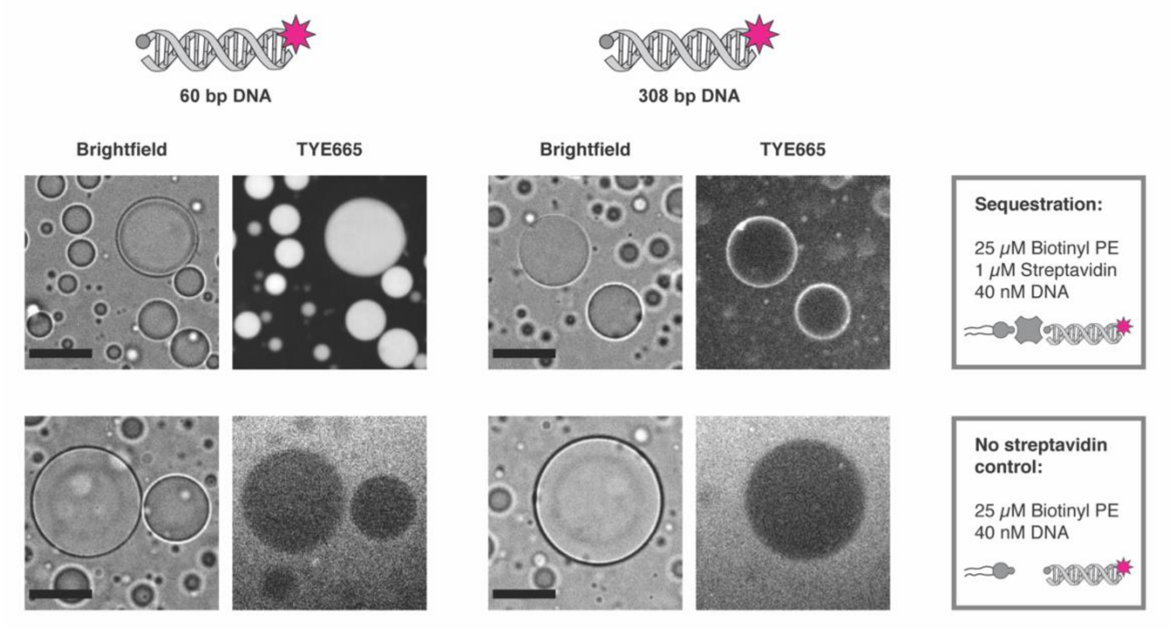


**Supplementary Fig. 2. Specificity of DNA sequestration.** Sequestration of biotinylated DNA requires the complete targeting system consisting of biotinyl PE, streptavidin and biotinylated DNA (top). In a control sample without streptavidin (bottom), short and long double-stranded DNA were excluded from the droplet phase and observed as diffuse fluorescence in the solution phase. Due to the high differences in fluorescence signals, intensities are not shown with the same brightness settings. Scale bars: 30 µm.

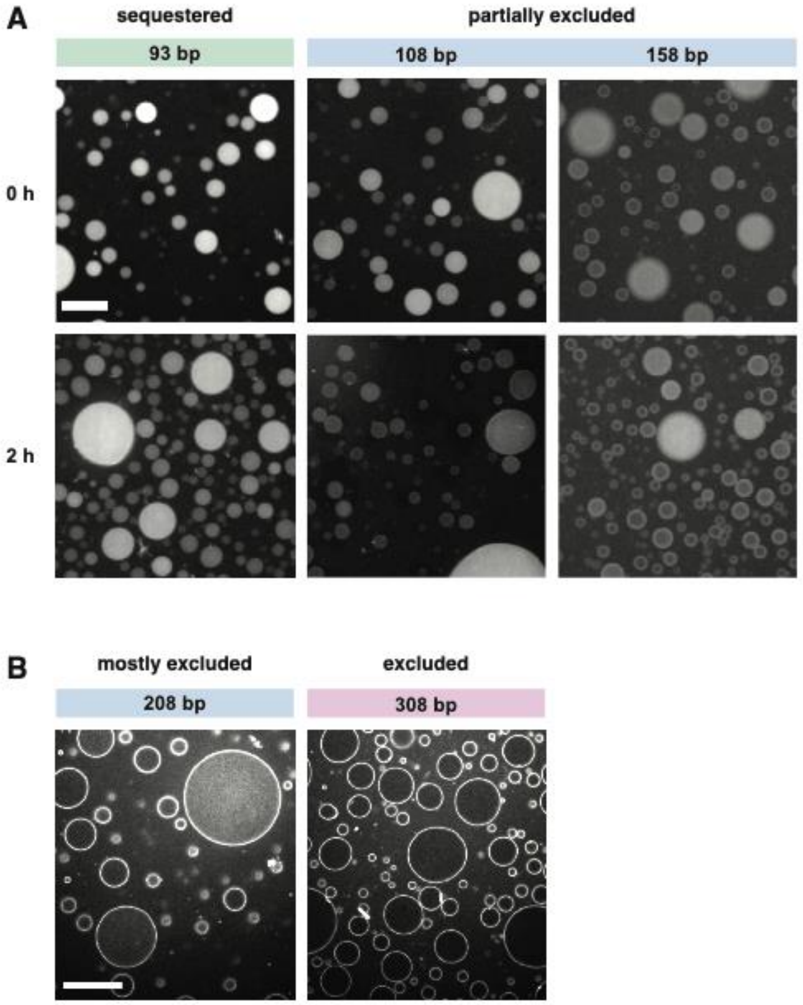


**Supplementary Fig. 3. Size-dependent transition from DNA sequestration to exclusion.** (**A**) The transition from sequestered to partially excluded states is observed as decrease in droplet fluorescence and the appearance of a bright corona around droplets that intensifies with time after mixing, particularly in small droplets. Images are shown at the same scale and with the same brightness settings in a logarithmic scale and were acquired with a 20x objective. (**B**) Transition from mostly excluded to fully excluded states imaged with a 63x objective. Images were acquired 2 h after sample preparation. Scale bars: 25 µm.

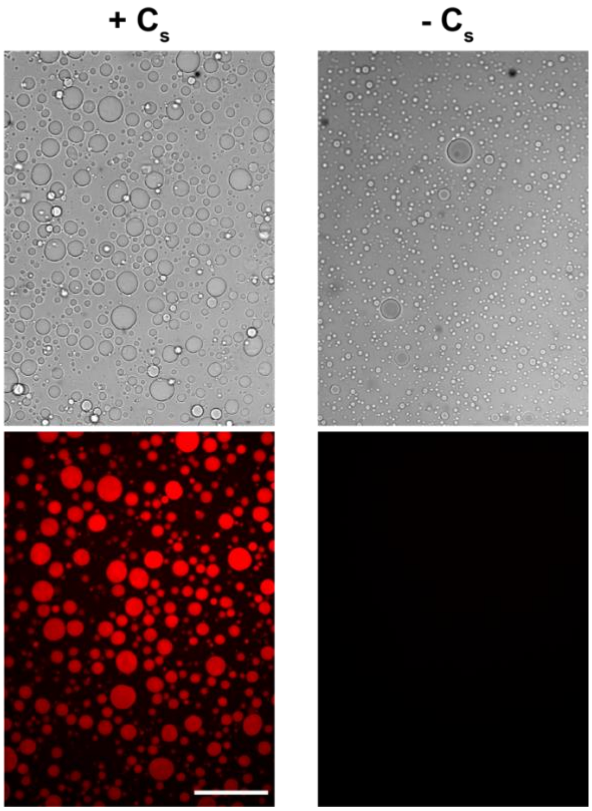


**Supplementary Fig. 4. Specific sequestration of $F_{red}$ DNA strand by hybridization to droplet-bound capture strand $C_s$.** Confocal microscopy images demonstrating $C_s$-mediated hybridization of fluorescently labeled $F_{red}$ (Top: bright field, bottom: TYE665 fluorescence). Fluorescence intensity increased only in droplets containing the capture strand $C_s$ (left). No changes in fluorescence intensity inside droplets prepared without $C_s$ (right). Scale bar: 100 μm.

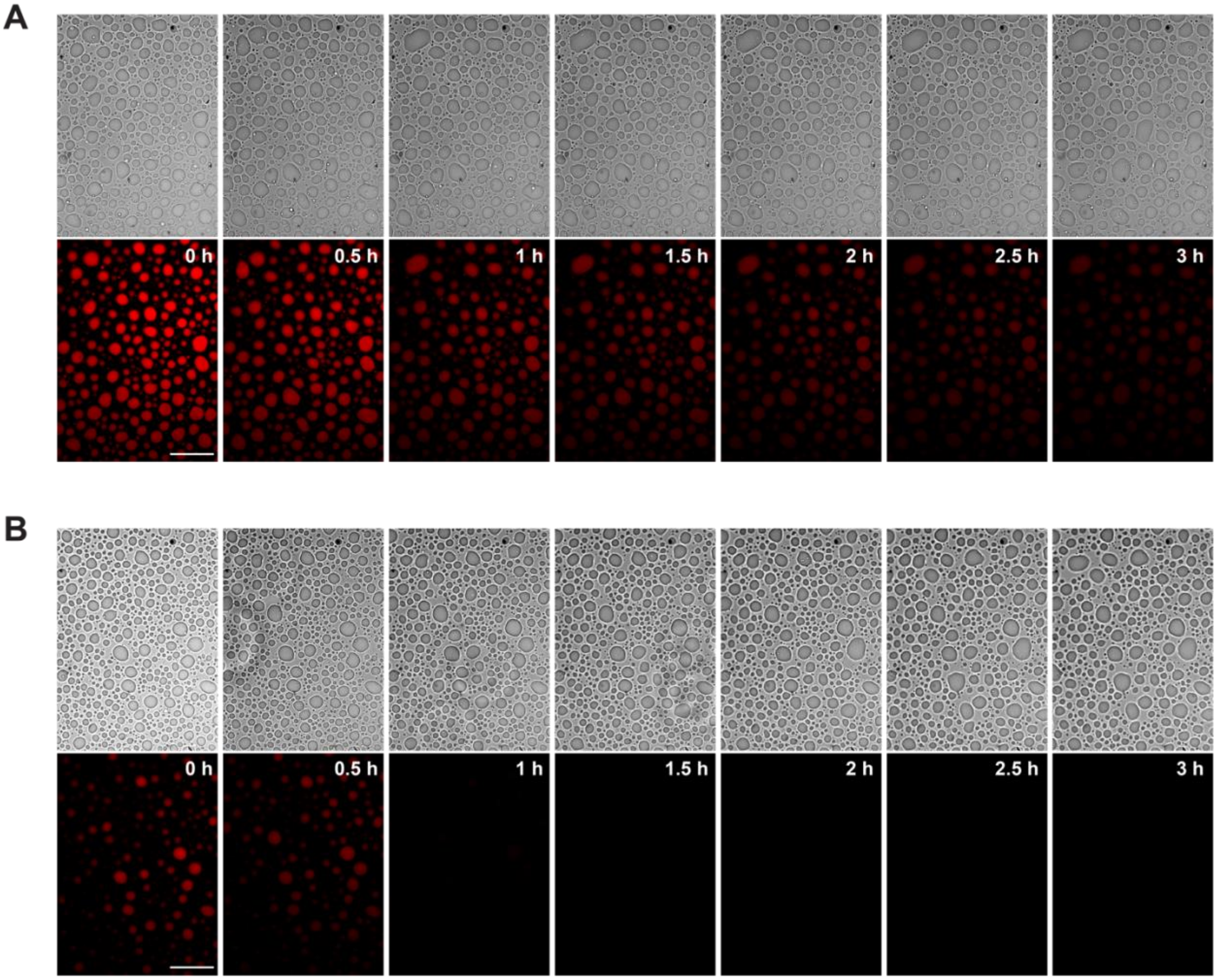


**Supplementary Fig. 5. Release kinetics.** Time-dependent confocal microscopy images showing toehold mediated displacement of $F_{red}$ by the addition of release strand F* on droplets containing the $C_d$ complex. (**A**) Decrease in droplet fluorescence upon displacement of $F_{red}$ (40 nM) with equimolar concentrations of F* (40 nM). (**B**) Accelerated release displacement of $F_{red}$ (40 nM) with two molar equivalents of F* (80 nM). $F_{red}$ was added at 0 h in both experiments. Scale bars: 100 μm.

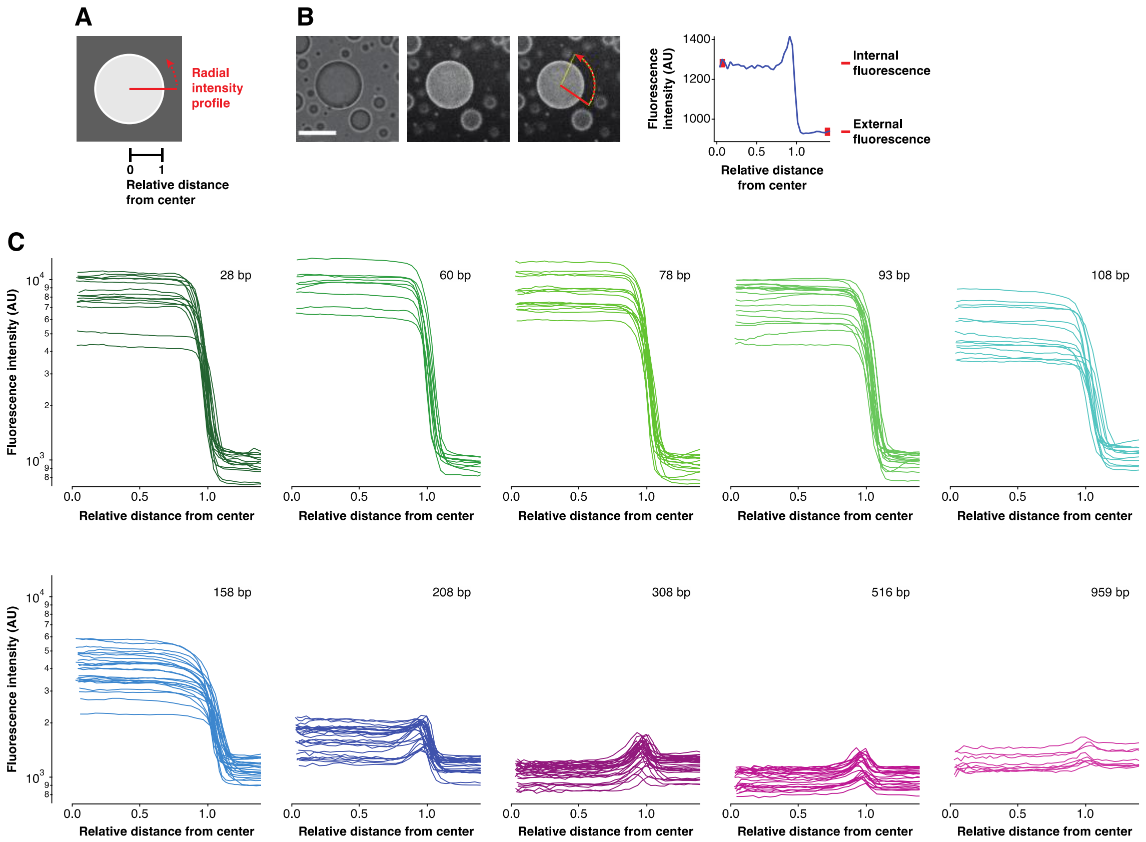


**Supplementary Fig. 6. Quantification of sequestration and exclusion by analysis of fluorescence intensity profiles of droplets.** (**A**) Schematic of the measurement of a radial intensity profile. (**B**) Example of a radial intensity profile measurement with Fiji/ImageJ as described in the methods. Shown are from left to right a brightfield image, corresponding TYE665 fluorescence image and the region that was used to measure the radial intensity profile. Note that the selected region does not contain any other fluorescent droplets so that it represents the fluorescence of the solution. The corresponding radial intensity profile is shown on the right. (**C**) Radial intensity profiles of all analyzed droplets for each DNA size (data in Fig. 2B).

**Supplementary Table 1.** Oligonucleotides for the PCR assembly of different length DNA constructs with TYE665 and biotin modifications.

| Size dsDNA (bp) | template or fwd primer name | Sequence (5′→3′) | rev primer name | Sequence (5′→3′) | assembly notes |
|---|---|---|---|---|---|
| 28 | rev_3'final_biotin | /5Biosg/GCTTGATATCGAATTCCTGCAGCCCGGG | 3'final-TYE | /5TYE665/cccgggctgcaggaattcgatatcaagc | annealed both oligos in buffer |
| 60 | 60bp | CGAGGATCTTAAGGCTAGAGggGCTTGATATCGAATTCCTGC | - | - | used 60bp oligo as template for PCR with final primers |
| 78 | 78bp | CGAGGATCTTAAGGCTAGAGggcatccgcttacagacaagGCTTGATATCGAATTCCTGC | - | - | used 78bp oligo as template for PCR with final primers |
| 93 | 93bp_1 | CGAGGATCTTAAGGCTAGAGggcatccgcttacagacaagctgtgac | 93bp_2 | GCAGGAATTCGATATCAAGCcggagacggtcacagcttgtctgtaagc | used mix of 93bp_1 and 93bp_2 oligo as template for PCR with final primers |
| 108 | fwd | CGAGGATCTTAAGGCTAGAGggcatccgcttacagacaag | 50-r | GCAGGAATTCGATATCAAGCgacacatgcagctcccggag | PCR on pET-Trx-mSa2 (Addgene #52320), then second PCR with final primers |
| 158 | fwd | CGAGGATCTTAAGGCTAGAGggcatccgcttacagacaag | 150-r | GCAGGAATTCGATATCAAGCgcggatgaacaggcagacat | PCR on pET-Trx-mSa2 (Addgene #52320), then second PCR with final primers |
| 208 | fwd | CGAGGATCTTAAGGCTAGAGggcatccgcttacagacaag | 250-r | GCAGGAATTCGATATCAAGCggaggcatcagtgaccaaac | PCR on pET-Trx-mSa2 (Addgene #52320), then second PCR with final primers |
| 308 | fwd | CGAGGATCTTAAGGCTAGAGggcatccgcttacagacaag | 458-r | GCAGGAATTCGATATCAAGCcctgtggaacacctacatct | PCR on pET-Trx-mSa2 (Addgene #52320), then second PCR with final primers |
| 516 | fwd | CGAGGATCTTAAGGCTAGAGggcatccgcttacagacaag | 700-r | GCAGGAATTCGATATCAAGCgcgcatgatcgtgctcctgt | PCR on pET-Trx-mSa2 (Addgene #52320), then second PCR with final primers |
| 959 | fwd | CGAGGATCTTAAGGCTAGAGggcatccgcttacagacaag | 901-r | GCAGGAATTCGATATCAAGCgcaactcgtaggacaggtgc | PCR on pET-Trx-mSa2 (Addgene #52320), then second PCR with final primers |
| all | 5'final-TYE | /5TYE665/gtcttcacctcgaggatcttaaggctagag | 3'final-biotin | /5Biosg/cccgggctgcaggaattcgatatcaagc | Final amplification of all DNA construct except 28 bp. |

**Supplementary Table 2.** Oligonucleotides for hybridization and toehold mediated strand displacement reaction in lipid sponge droplets.

| Name | Sequence (5′→3′) | Name | Sequence (5′→3′) |
|---|---|---|---|
| $C_s$ | /5Biosg/AAAACCCTCATTCAATACCCTACG | $F_{red}$ | /5TYE665/GAAGTGACATGGAGACGTAGGGTATTGAATGAGGG |
| F* | CCCTCATTCAATACCCTACGTCTCCATGTCACTTC | $F_{green}$ | /5ATTO488N/GAAGTGACATGGAGACGTAGGGTATTGAATGAGGG |
